# Immunological Diversity with Similarity

**DOI:** 10.1101/483131

**Authors:** Rohit Arora, Harry M. Burke, Ramy Arnaout

## Abstract

A diverse immune repertoire is considered a hallmark of good health, but measuring diversity requires a framework that incorporates not only sequences’ relative frequencies but also their functional similarity to each other. Using experimentally measured dissociation constants from over 1,300 antibody-antigen and T-cell receptor (TCR)-peptide pairs, we developed a framework for functional immunological diversity based on binding and applied it to nearly 400 high-throughput antibody and TCR repertoires to reveal patterns in immunological memory, infection, vaccination, and aging. We show that functional diversity adds information that is not captured by raw diversity, revealing signatures of e.g. clonal selection, and that unlike raw diversity, functional diversity is a robust measure that does not require correction for sampling error. Finally, we show that according to functional diversity, unlike raw diversity, individuals’ repertoires overlap substantially, indicating a definable ceiling for the functional diversity of human adaptive immunity. Similarity redefines diversity in complex systems.

---

Immune repertoires are famously diverse. Collectively, a person’s ~10^12^ B and T cells express many millions of unique recombined antibody and TCR genes as part of millions of clonal lineages, more unique sequence than in the entire germline genome^1^. At the sequence level, the repertoires of any two people overlap by only a fraction of a percent, indicating still higher diversity in the population^2,3^. Yet repertoires are formed from V, D, and J gene segments that almost all people share and that are expressed at similar frequencies across individuals, and most repertoires are shaped by similar antigenic exposures and a consequent need to recognize and bind similar targets^4,5^. Squaring the diversity that is seen with the similarity that must exist is a major goal of high-throughput immunology.

This goal has relevance for disease stratification and clinical management across a range of conditions. B- and T-cell diversity fall with age, as specific exposures expand a few select lineages at the expense of others^6^. Chronic infection appears to have a similar effect, impairing vaccination^1^. Low B-cell diversity is associated with physiological frailty, a syndrome seen alongside conditions that have traditionally been considered to be unrelated to adaptive immunity (e.g., atherosclerotic cardiovascular disease), independent of chronological age^7^. In cancer, a rise in sequence-level T-cell diversity is thought to predict a successful response to immune-checkpoint inhibitors, drugs that make tumors more visible to the immune system^8^.

Two key contributors to any intuitive measure of diversity are frequency and similarity, but traditionally only frequency has factored into standard measures of immunological diversity. The simplest measure, species richness—the number of different sequences, lineages, or clones in a sample—ignores frequency entirely. Yet in a practical sense, a repertoire with a single dominant (e.g., leukemic) clone should be considered less diverse than a repertoire that has the same number of clones but no dominant clone. To incorporate frequency into measurements of diversity, there exist a family of measures—species richness (as a limiting case), Shannon entropy, the Simpson index (and related Gini coefficient), and the Berger-Parker index, among others—that differ in how much weight each measure places on frequency: i.e., in how much more a large clone adds to the total diversity than a small one^9^. Mathematically, this family is derived from the single master equation

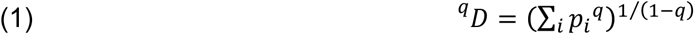

in which the different measures correspond to differences in the “frequency parameter” *q* (e.g. *q*=0 corresponds to species richness; *q*=1 to Shannon entropy), ^*q*^*D* is read “*D*-*q*” (“diversity of order *q*”), and *i* indexes the different species present in the sample. No single diversity measure is “best:” the different measures are complementary single-parameter summaries of the underlying species-frequency distribution^10^. ^*q*^*D* for immunological diversity is subject to considerable sampling error, but robust methods exist for correcting for this error and are becoming standard in the field^11,12^.

In the present work our goal was to incorporate and assess the utility of the second key contributor: species similarity. A repertoire made up of all-different sequences is intuitively more diverse than a repertoire that has the same number of sequences, present at the same frequencies as in the first repertoire, but all drawn from the same lineage or clone. Although our focus here is immunological diversity, similarity is a key contributor to diversity in many complex systems, including metagenomics (species vs. operational taxonomic units), machine learning (similarities within training data vs. the size of the training set), and transcriptomics (cell-to-cell variation vs. cell types; see Discussion). In the immune-repertoire literature, this point is sometimes addressed indirectly by grouping sequences together before measuring diversity, for example by clustering reads, collapsing clones, or binning by V(D)J segment usage^1,5,13–15^. However, grouping usually imposes a binary threshold—in or out—on what is by nature a continuous and overlapping relationship among sequences and their encoded proteins. Grouping also usually zeros out or ignores any diversity that might exist within groups. It is unclear what is lost by ignoring similarity, or what might be gained from a more complete synthesis of diversity with similarity. Therefore we sought to develop and explore a continuous framework for immunological diversity incorporating similarity, and to test its utility in situations of clinical and research interest.

## Results

### Framework

We measured diversity-with-similarity on high-throughput B- and T-cell repertoires using a robust mathematical framework initially proposed for studying diversity in ecology and environmental settings^16^:

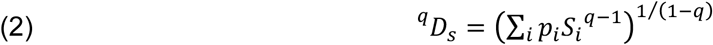

where

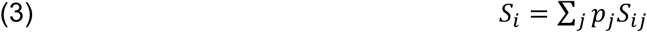

Here the key innovation relative to Eq. 1 is a similarity matrix *S*, whose entries *S*_*ij*_ quantify how similar each pair of species (sequences, clonotypes, etc.) is. This framework provides “with-similarity” counterparts for species richness and the frequency-weighted diversity measures: species richness with similarity (^0^*D*_*s*_, which places a very small weight on frequency, and ^∅^*D*_*s*_, which like ^0^*D* ignores frequency altogether), the exponential form of entropy with similarity (^1^*D*_*s*_, henceforth simply “entropy with similarity,” and likewise for other named measures), and so on. In ^*q*^*D*_*s*_ notation, *q* as before is the frequency parameter, *D*_*s*_ denotes diversity-with-similarity, and *D* without the subscript means diversity without similarity, which we refer to as “raw diversity” (see Methods for more on notation).

Constructing the similarity matrix necessitates a choice of similarity measure. (Note the difference between similarity measures and diversity measures: a similarity measure is used to build the similarity matrix, which then is used to calculate diversity measures.) The choice of similarity measure is up to the investigator and depends on the biological feature(s) of interest. For example to study somatic hypermutation, one might use the Hamming distance. Just as different weightings provide complementary information about rare vs. frequent species—for example, the number of new thymic emigrants (species richness; ^0^*D* or ^∅^*D*_*s*_) vs. large leukemic clones (Berger-Parker index; ^∞^*D*)—different similarity measures are expected to reveal different systems-level features of repertoires’ sequence-level configuration. Also as with raw diversity measures, expressing results for the new diversity-with-similarity measures as effective numbers, also known as number equivalents^9,17–19^, as opposed to as bits or nats (for ^1^*D*_*s*_) or as various fractions (for ^>1^*D*_*s*_), makes it possible to compare them to each other, regardless of weighting or similarity measure, on a single intuitive scale (Box 1).

##### Box 1: Glossary

###### Diversity

Any of several measures of the number of species present in a population, possibly accounting for the relative frequencies of, or pairwise similarity between, species (e.g. species richness, Shannon entropy, Gini coefficient, Berger-Parker index)

###### Frequency

The relative number of each species in the population under study, which is taken into account by most measures of diversity but not species richness (which is a simple count)

###### Similarity

An investigator-defined measure of how alike or different a given pair of species are to each other (e.g. the Hamming distance between each pair of sequences)

###### Effective numbers

A method for comparing different diversity measures in the same units, i.e. numbers of species. Consider two repertoires of 100 clones each. In the first repertoire, one clone is large and accounts for 91 percent of all cells (e.g. a leukemic clone); the other 99 clones are small and account for the remaining 9 percent. In the second repertoire, all 100 clones are equally common, each accounting for 1 percent of cells. The Shannon entropies of the two repertoires are 1.0 bit and 6.6 bits. Entropy is converted to an effective number—^1^*D*—by exponentiation: the effective number of clones in the first repertoire is 2^1.0^=2, while in the second repertoire it is 2^6.6^=100. Thus per entropy, the first repertoire can be thought of as “effectively” consisting of just two clones: the 99 rare clones collectively count the same as the one large clone. In other words, the first repertoire has the same effective diversity as a repertoire that consists of just two clones that are equally common.

The second repertoire already consists of clones that are equally common, so the effective number of clones in this repertoire, 2^6.6^=100, is the same as its species richness. Diversity with similarity is interpreted in a similar fashion: a repertoire with a ^*q*^*D*_*s*_ of *n* species has the same effective diversity as a repertoire with *n* species that are equally common (as above), with the additional constraint that these species are now also completely unrelated to/dissimilar from each other.

### Similarity measure

We were interested in the single most fundamental mechanistic feature of antibodies and TCRs: binding to specific targets (Fig. 1). Therefore for our similarity measure, we chose a proxy for binding affinity that follows from the empirically observed changes in dissociation constant (*K*_*d*_) associated with amino-acid substitution in antibody and TCR CDRs^20^. We found that on average, a single amino-acid substitution at an antibody-antigen or TCR-peptide binding surface lowers affinity by 4-5 fold (geometric mean), with a long tail corresponding to rare orders-of-magnitude effects (Fig. 2a). We focused on CDR3, the third complementarity determining region, of IgH and TCRβ, since this is the single most important contributor to binding specificity^21^; however, our approach can be applied to other regions (or indeed other protein families). Because the relationship between sequence and specificity remains non-predictive and therefore complex, for any given sequence pair the similarity imputed from the observed distribution will be approximate; however, averaged over the many millions of pairs in the average high-throughput repertoire, it was expected to be a reasonably accurate first-pass repertoire-level view of immunological diversity with binding similarity. Using this similarity measure, diversity-with-similarity is interpreted as the effective number of sequences in a repertoire if the sequences were equally common and had no binding overlap with each other (Box 1), or equivalently, the number of equally common non-overlapping binding targets that a repertoire can recognize. We therefore refer to this choice of diversity-with-similarity as “functional diversity” (Fig. 1). Functional diversity can be interpreted in the context of a “shape space” ^22^ that contains all possible CDR3 binding targets, with nearby targets having similar three-dimensional shapes and conformations (Fig. 1a). Each CDR3 binds a (possibly overlapping) subset of targets; together, a repertoire's CDR3s cover some part of shape space (Fig. 1b). Functional diversity measures the size of this region, controlling for similarity and overlap in binding among different CDR3s (Fig. 1c).

**Figure 1:**
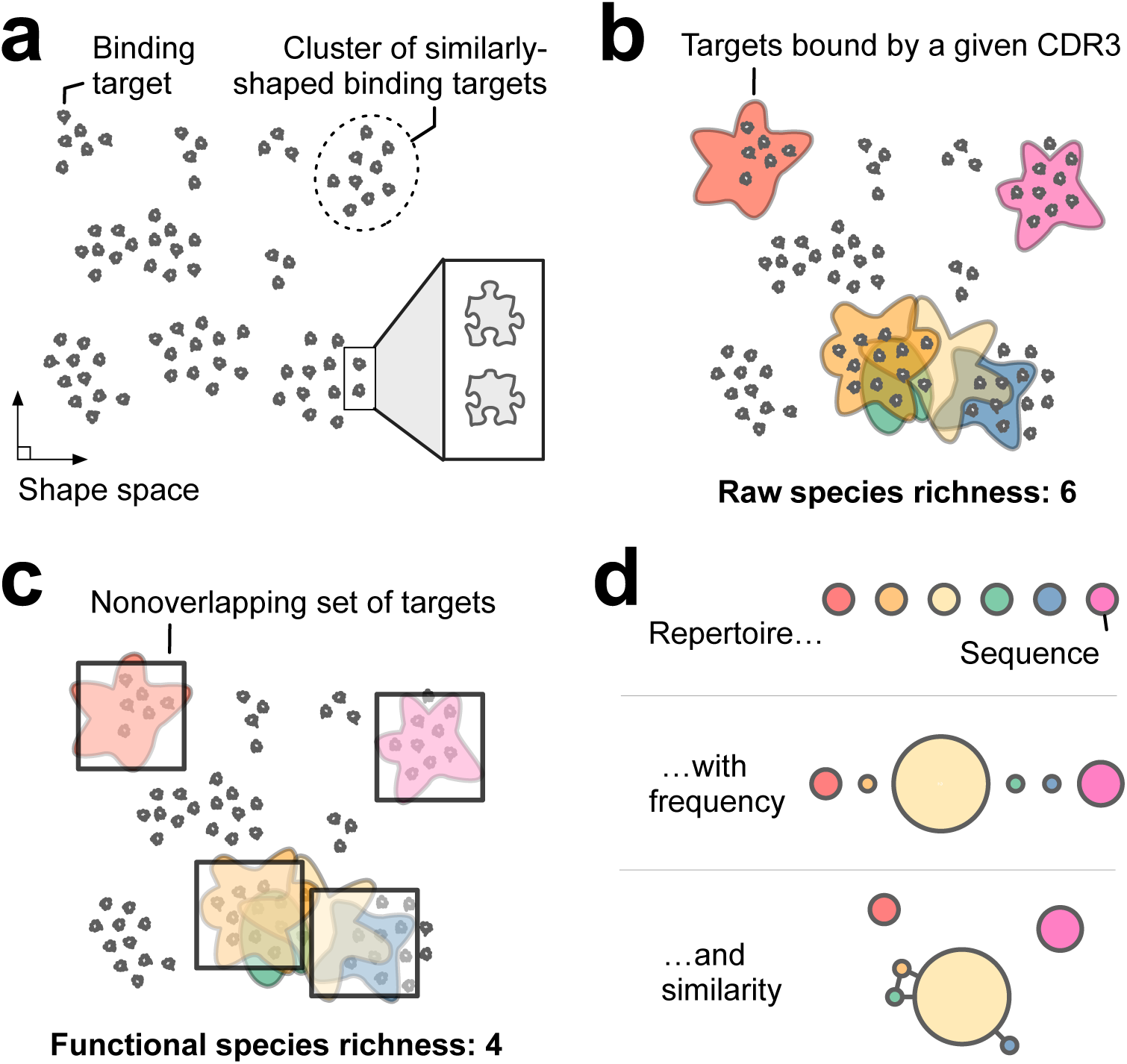
Functional diversity. **(a)** Each dot represents a binding target (e.g. an epitope) with a different shape. Nearby targets have similar shapes (inset). Targets form clusters of similarity. **(b)** Each colored region represents the targets that can be bound by one of six unique CDR3 sequences in a representative repertoire; this repertoire has a raw species richness of 6. Together, the colored regions cover the part of shape space that can be bound by the repertoire. (The unbound region might include, e.g., self-antigens.) Note the substantial overlap in binding targets for the orange, yellow, green, and blue CDR3s. This overlap reflects binding similarity among these CDR3s. **(c)** Because of this similarity, the repertoire covers only the region denoted by the four identical non-overlapping squares. The functional species richness of this repertoire is therefore 4: this repertoire has the same species richness as a repertoire comprising four CDR3s that have zero overlap in binding specificity. **(d)** Schematic representation of repertoires without incorporating frequency or similarity (top), with frequency only (middle), and with frequency and similarity (bottom).

**Figure 2:**
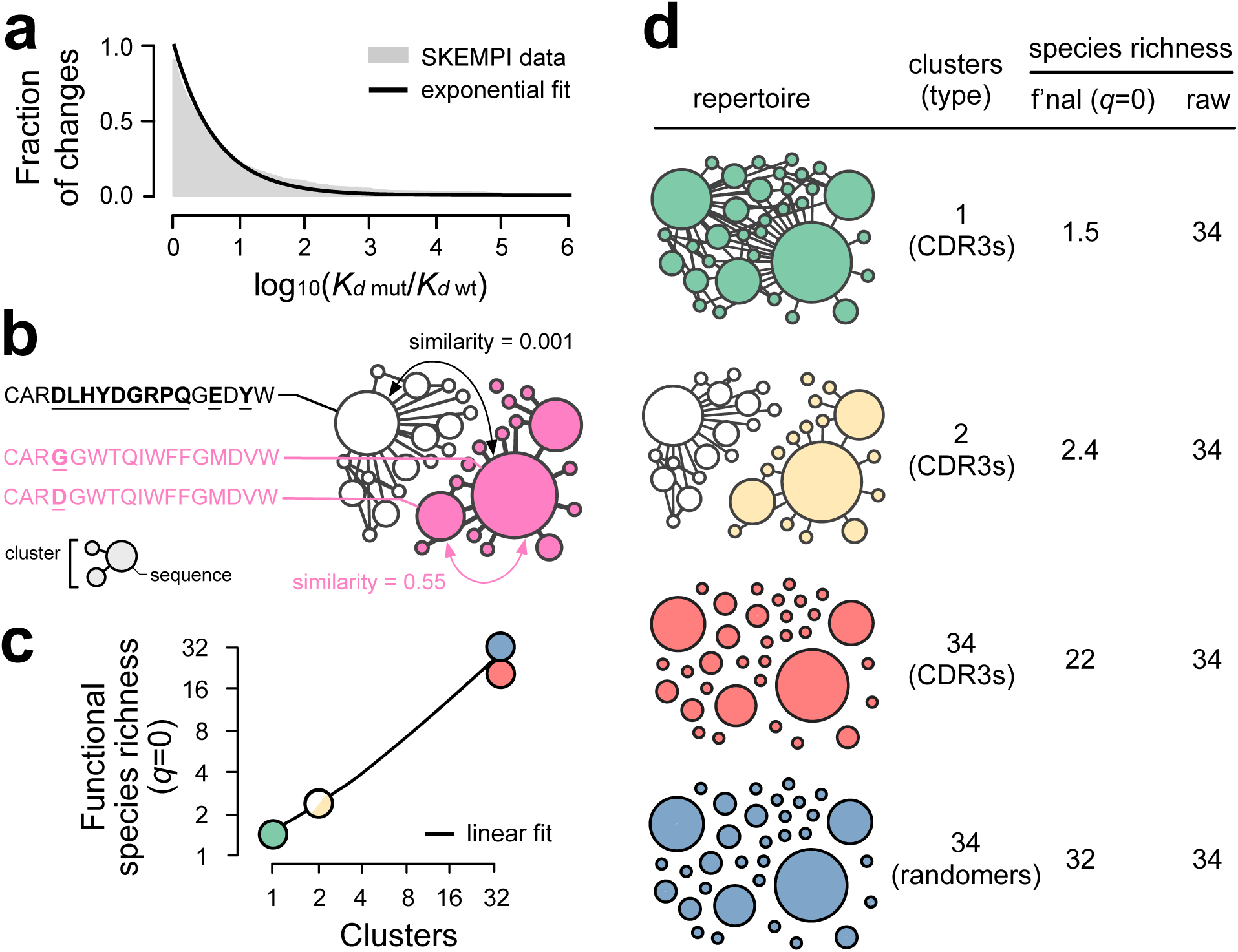
Validity. **(a)** Single amino-acid mutations in antibody and TCR molecules have a range of effects on affinity, as measured by change in dissociation constant, *K*_*d*_ (gray). This was well fit by a simple exponential (black line), providing parameterization for the similarity metric. **(b)** CDR3s with high sequence identity have high similarity, while different CDR3s have low similarity. Shown are two clones, represented by red and white subnetworks, each composed of 17 unique CDR3 sequences drawn from clonotypes of two real IgH repertoires. Node size corresponds to the frequency of each sequence; edges connect pairs of sequences that differ at a single amino-acid position. **(c)-(d)** Functional similarity agrees with an intuitive sense of repertoire diversity. Each of the four repertoires in (c) has the same number of unique sequences, present at the same frequencies; as a result, they all have identical raw diversity (for every value of *q*) despite their obvious quantitative and qualitative differences. In contrast, functional diversity increases with the number of, and increasing difference between, repertoires' constituent sequences. Node size denotes sequence frequency. Shades denote different clonotypes. Comparing the third and fourth rows, note that even when two repertoires have the same number and frequencies of unique sequences, the repertoire whose sequences are more different from each other (random peptides) has the higher functional diversity.

### Validity

We first established that our similarity measure behaved sensibly by testing that closely related sequences scored close to 1 and unrelated sequences scored close to zero (in a continuous manner, following from an assumption of multiplicative independence and the data in Fig. 2a; see Methods) (Fig. 2b). We next established that it resulted in intuitive values for functional similarity by testing against expectations on simple *in silico* repertoires. In a representative test, we constructed four repertoires with 34 unique sequences each and 752 sequences total (Fig. 2c-d). In each repertoire, a few sequences were common (larger circles) while most were rare (smaller circles), representing the long-tailed frequency distribution seen in real repertoires^3,23^. Importantly, the species-frequency distribution for all four repertoires was identical, meaning that raw diversity was also identical across the repertoires, for all frequency weightings. The only difference between the repertoires was in the pairwise similarity among sequences.

For the first repertoire in this illustrative example (Fig. 2d, top row), we chose closely related sequences from a single real-world CDR3 clone. We expected that species richness with similarity—functional species richness—would be close to 1. (We used ^0^*D*_*s*_ here; ^∅^*D*_*s*_ performed similarly.) We observed a value of 1.5; the extra 0.5 reflected sequence diversity within the clone. For the second repertoire (Fig. 2d, second row), we swapped out half the unique CDR3s with CDR3s from a different, unrelated real-world clone. As expected, we observed a rough doubling of functional diversity, to 2.4. For the third repertoire (Fig. 2d, third row), we replaced all the sequences with 34 randomly chosen real-world CDR3s. We expected a functional diversity that was much higher than in the first two repertoires but less than 34 because of the inherent sequence similarities that make a CDR3 a CDR3, and, consistent with this expectation, observed a value of 22. For the final repertoire (Fig 2d, bottom row), we replaced the CDR3s with random amino-acid sequences (controlling for length), expecting a functional similarity of nearly 34, and this again was observed (^0^*D*_*s*_=32). In contrast to these differences in functional diversity, raw diversity was indistinguishably 34 for all four repertoires. In every example, functional diversity fit an intuitive sense of what diversity should mean (Fig. 2c), while raw diversity failed to detect the defining differences among these repertoires. The agreement between expectation and observation supports the validity of our functional-diversity framework for immune repertoires.

### Robustness

Sampling error—the “missing-species” problem^24^—is known to be a major potential confounder when measuring raw diversity, necessitating large (e.g.) blood volumes and/or post-hoc statistical correction for measurements on the sample to reflect repertoire diversity in the individual as a whole^11^. A practical feature of functional similarity is that the values are smaller than those of raw diversity (reflecting clustering of similar sequences; Fig. 1-2). The effective coverage is therefore greater, meaning that less information about the functional diversity of an individual’s overall repertoire is lost upon sampling than is the case for raw diversity. This observation suggested that functional diversity is more robust to sampling error, possibly even making it accurate enough to use without statistical correction, and thus useful for the sample sizes typically available for sequencing (10,000-1 million cells).

To test this possibility, we systematically downsampled from a representative TCRβ repertoire and two representative IgH (IgG) repertoires, one prepared from mRNA and one from genomic DNA, each with ~10^6^ unique sequences, and compared raw vs. functional diversity on the subsamples to those of the full sample (Fig. 3). (We tested mRNA and DNA separately to test the possibility of lower diversity from mRNA than DNA, which was expected since transcriptionally less-active cells may be less likely to be sampled.) For TCRβ, we found that functional species richness saturated at a sample size of ~30,000 sequences and functional entropy at ~10,000 sequences (Fig. 3b, first and third columns). Functional diversity for higher frequency-parameter values (*q*) saturated with even fewer sequences. For IgH from mRNA, functional species richness did not saturate but did plateau, with a final increase of ≤2 percent per order of magnitude. Assuming that each unique sequence corresponds to a cell and 10^11^ B cells in the body, this final measured rate of increase means that the individual's total functional species richness is no more than 50 percent higher than the value measured on the sample (Fig. 3, middle row). This is the maximum expected sampling error. For IgH from DNA, functional species richness had begun to plateau at the full sample size, resulting in the value for the individual being no more than three times as much as in the sample (maximum three-fold error). Meanwhile, functional entropy saturated at 30,000 cells for IgH from mRNA and 300,000 cells for IgH from DNA. This behavior was in marked contrast to that of raw diversity, which did not saturate or plateau for species richness (Fig. 3, white symbols), consistent with previous reports and illustrating the need for statistical correction^11^.

**Figure 3:**
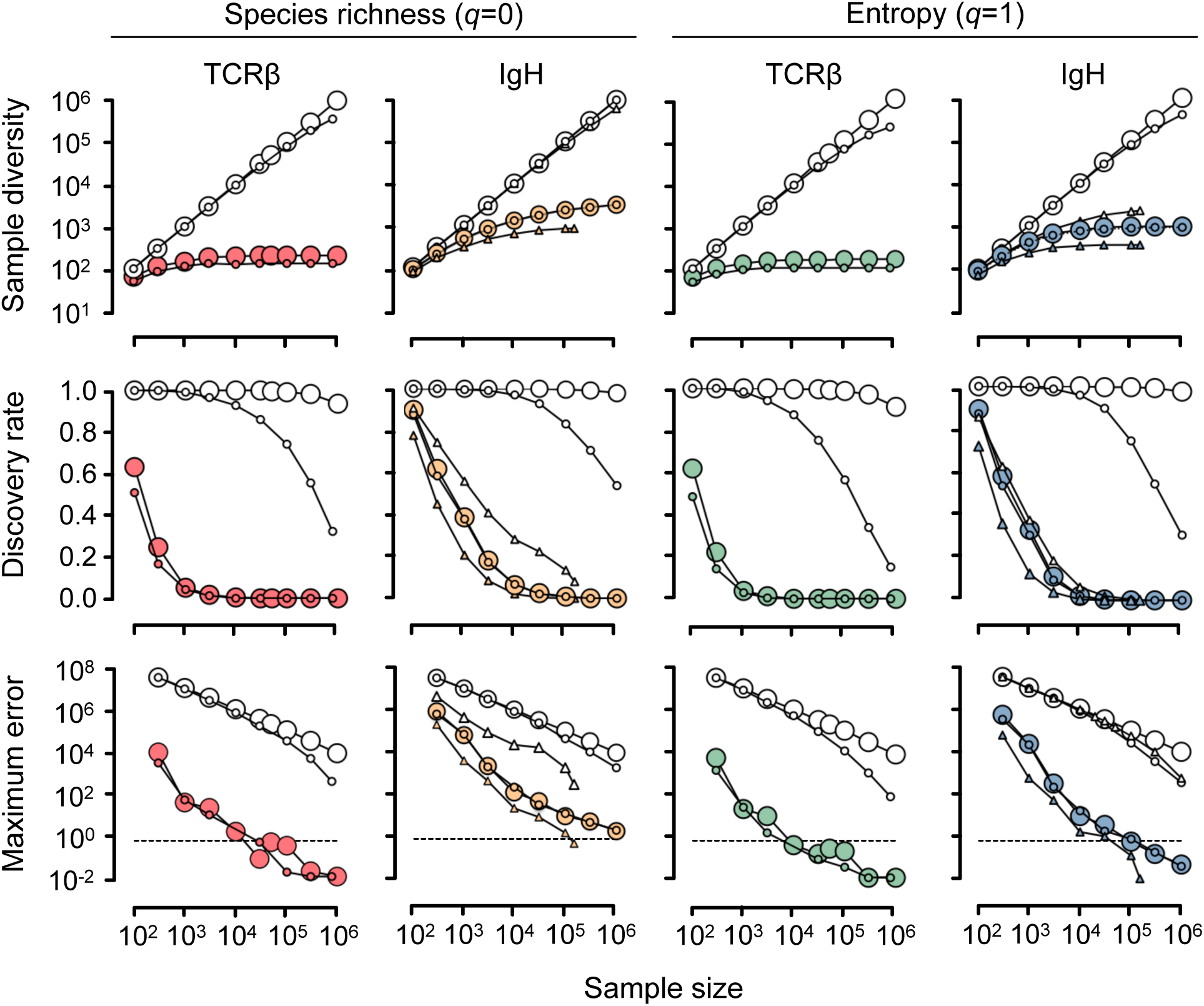
Robustness. Results for raw and functional species richness (*q*=0; ^0^*D* and ^0^*D*_*s*_) and raw and functional entropy (*q*=1; ^1^*D* and ^1^*D*_*s*_). Raw, white shapes; functional, colored shapes. Large symbols give an upper bound/worst-case scenario based on sampling meta-repertoires; small symbols give results for a representative sample from DNA (circles) and mRNA (triangles). First row: sample diversity is plotted as the effective number of sequences. For *q*=0, functional diversity plateaus for TCRβ and IgH RNA and trends toward a plateau for IgH DNA at the tested sample sizes; all three plateau for *q*=1; raw diversity does not plateau for either *q*=0 or *q*=1. Second row: discovery rate is the probability that the next sampled sequence will add to the diversity. For example, for the IgH DNA representative sample, for *q*=0 raw diversity, at a sample size of 1 million sequences there is still a probability of ~0.5 (a 50 percent chance) that the next sequence to be sampled will be one that has not yet been seen and will therefore add to the diversity. Third row: maximum error is the maximum fraction by which the diversity in the sample can underestimate the diversity in the individual from whom the sample was taken. Horizontal dashed lines indicate the threshold for two-fold error. For example, for the worst-case scenario for TCRβ, *q*=0 functional diversity measured on a sample of 10,000 sequences will be no more than a two-fold underestimate of diversity in the individual as a whole; in other words, the sample value will be at least 50 percent of the overall value.

We then asked whether the robustness of functional diversity can be expected to generalize for any IgH or TCR repertoire. We reasoned that a “meta-repertoire” comprising sequences drawn uniformly (i.e., without regard to frequency) from many individuals will be more diverse, by any measure, than any single repertoire (which will have fewer sequences, and in which the same sequence may appear multiple times, resulting in lower diversity for *q*≥0). Downsampling from a meta-repertoire therefore provides an upper bound or worst-case scenario for the sampling requirements for any single repertoire. To build meta-repertoires, we pooled and then uniqued CDR3s from 114 different IgH repertoires from 79 individuals including Americans of African, European, and Hispanic descent^5,15,25^ to build an IgH meta-repertoire of roughly 36 million unique sequences—as many as or more than ever observed or currently estimated to be in a typical individual—and similarly for CDR3s from TCRβ repertoires from 69 healthy individuals (of mostly European but some Asian descent)^**26**^ to build a TCRβ meta-repertoire of 10 million unique sequences, and downsampled from each of these meta-repertoires as above (Fig. 3, large circles). We chose healthy individuals to avoid any down-biasing of diversity that might come with sampling many related sequences e.g. from expanded somatically hypermutated clones (for IgH). We found that functional diversity plateaued for all *q*, saturated for *q*≥1 and reflected overall diversity to within a few percent from sample sizes of 50,000 for TCRβ and IgH RNA and 100,000 for IgH DNA for *q*=0, and 30,000 for TCRβ and IgH RNA and 300,000 for IgH DNA for *q*≥1 (Fig. 3, large colored circles). Together, these results confirmed that functional diversity measured on samples is an accurate measure of overall functional diversity in the individual, at conventional sample sizes.

### Raw and functional diversity

We measured raw and functional diversity on 141 healthy human subjects (Fig. 4). For IgH, we found a (geometric) mean functional species richness (^∅^*D*_*s*_) of 677 (range, 487-916) from mRNA and 2,205 (range, 2,042-2,485) from DNA, suggesting that on average, the human antibody repertoire is capable of recognizing the equivalent of no more than a few thousand unique non-overlapping heavy-chain CDR3 binding targets. (As above, lower diversity from mRNA was not unexpected, since inactive cells, which produce less IgH mRNA than active cells, may be underrepresented.) For TCRβ the mean functional diversity was 140 targets (range, 115-167). Functional diversity can be thought of as clustering similar sequences together, although functional clusters can overlap and sequences can belong to multiple clusters. An indication of the average size of these clusters can be obtained by taking the ratio of raw to functional diversity measures. For species richness, we found that IgH typically had on average hundreds of sequences per cluster, while TCRβ had thousands. Thus by both functional diversity and average functional-cluster size, IgH CDR3 repertoires are roughly 5-10 times as diverse as TCRβ (for small *q*). Repertoires with higher raw diversity might be expected to be more functionally diverse, but we found no consistent trend across all repertoire types (measured by correlation coefficient). Thus, functional diversity generally complements raw diversity, adding information not captured by raw diversity alone.

**Figure 4:**
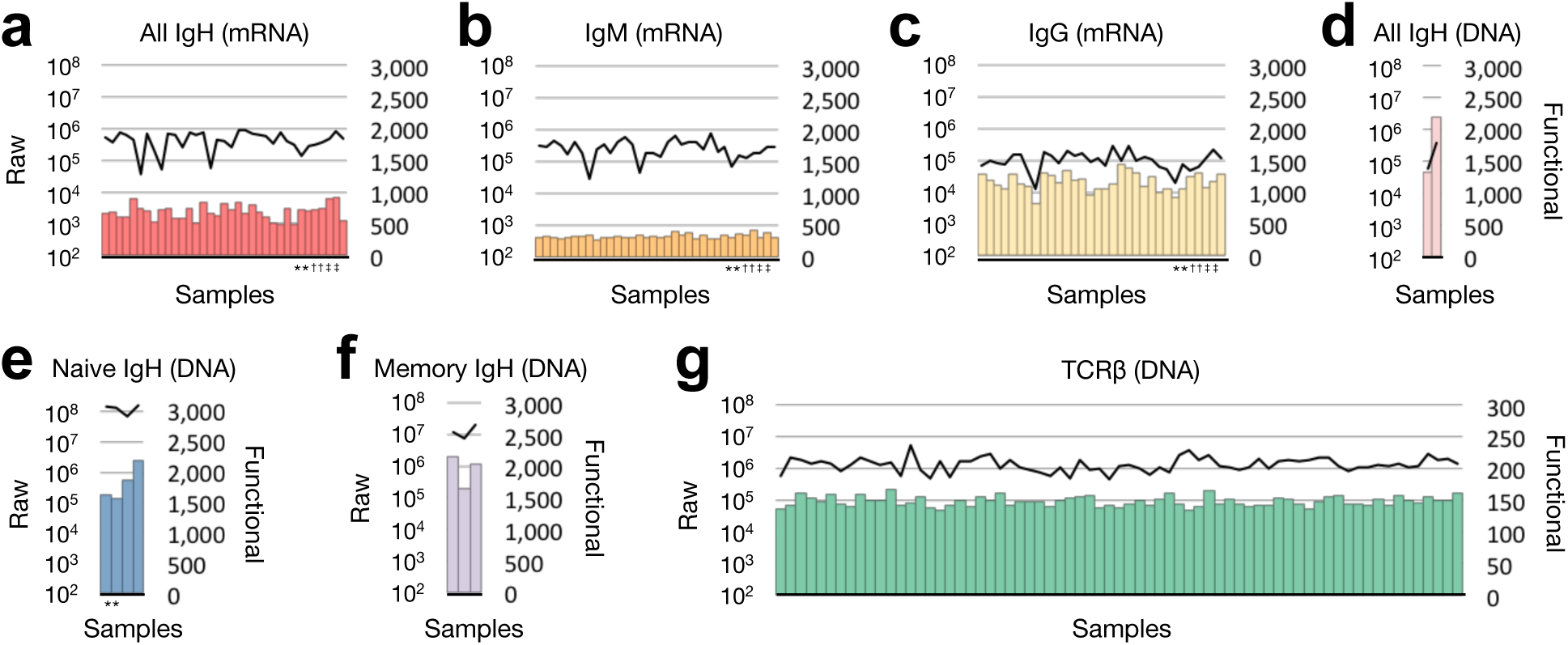
Diversity in individuals. Raw (black lines; left vertical axis) and functional (colored bars; right vertical axis) species richness (q=∅) for 179 CDR3 repertoires from healthy individuals representing **(a)** IgH from mRNA (all isotypes: IgA, IgG, IgM, IgD, and IgE), **(b)** IgM and **(c)** IgG from mRNA from the subjects in (a), **(d)** IgH from DNA (all isotypes), **(e)** naïve IgH from DNA, **(f)** memory IgH from DNA, and **(g)** TCRβ from DNA. See Methods for references. Matched pairs of symbols below the horizontal axis denote replicates. Note the difference in the scale for functional diversity between IgH and TCRβ. Note also a general lack of correlation between raw and functional species richness, except in (c).

### Naïve vs. memory B cells

We next sought to investigate what the complementary information provided by functional diversity might add to our understanding of adaptive immunity. We began by investigating two widely studied B-cell subsets, naïve (IgM) and memory cells (predominantly IgG). Previous studies have shown that naïve repertoires have higher raw diversity than memory repertoires^5^. This is at least superficially consistent with the fact that only a subset of naïve cells are selected to enter into the memory compartment. However, in a functional sense there is a case to be made that memory/IgG repertoires should be considered more diverse, since somatic hypermutation differentiates and thereby distances memory cells from naïve cells, and indeed from each other. Using well-characterized publicly available repertoires from DNA from three healthy human subjects, we confirmed that by raw species richness, naïve (CD27^−^IgM^+^) B-cell repertoires are ~10 times as diverse as memory (CD27^+^IgM^−^) repertoires ^7^ (Fig. 5a-b). Yet by functional species richness, we found that memory repertoires were at least as diverse as naïve (Figs. 5a-b). Comparing raw and functional diversity for 34 IgM and 32 IgG repertoires from mRNA (repertoires with less than 100,000 total sequences were discarded) from 28 additional healthy individuals from a separate dataset showed a similar pattern as for the three DNA repertoires: in all but a few outliers, IgM had higher raw diversity but IgG had higher functional diversity (Fig. 5c-d). For raw diversity, the IgM:IgG ratio rose from ~3:1 at *q*=0 to peak at 10:1 around *q*=1, due to a large fraction of rare IgG sequences (Fig. 5d). This effect was more pronounced for the naïve:memory ratio in the DNA dataset (Fig. 5b). For functional diversity, the absence of a peak in the IgM:IgG ratio suggests that these many rare sequences must nonetheless be similar to others in the repertoire, possibly because they are clonally related (Fig. 5b,d).

**Figure 5:**
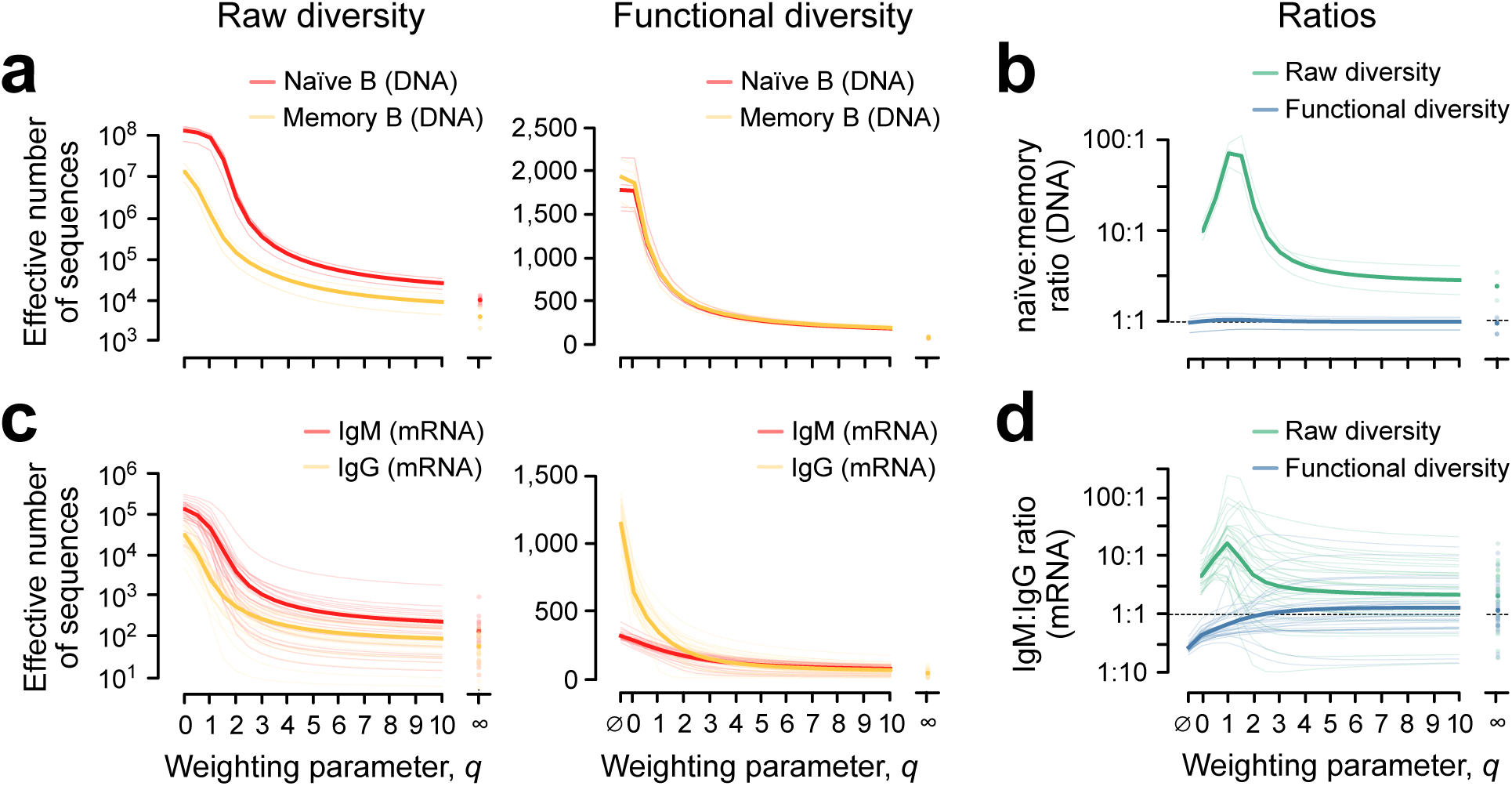
Naïve vs. memory. **(a)** Diversity profiles for naïve (red) and memory (black) compartments from three deeply sequenced individuals. A diversity profile is a way to show diversity across a range frequency-weighting parameter values at once. By raw diversity (left), the naïve compartment is more diverse across the range of weightings. By functional diversity (right), this distinction disappears. In **(b)**, this disappearance is highlighted by plotting the ratio of naïve:memory diversity for raw diversity (red) and functional diversity (black). According to functional diversity, the naïve compartment is no more diverse, and indeed sometimes somewhat less diverse, than the memory compartment. This reversal is even more prominent in comparisons of repertoires from an additional 28 healthy subjects **(c,d)**.

### Cytomegalovirus (CMV) exposure

CMV is a herpesvirus to which half of the adult population has been exposed and results in life-threatening opportunistic infections in newborns, transplant recipients, and immunocompromised individuals^27^. In most healthy individuals, it causes a chronic infection marked by clonal expansion of both B cells and T cells and a consequent fall in raw diversity, an effect also seen during aging (see below)^4,28^. We measured raw and functional TCRβ CDR3 diversity for 120 individuals: 69 CMV-seronegative and 51 CMV-seropositive subjects aged 19-35^26^, the narrow age range helping control for any age-related effects. There was a clear trend toward lower diversity in the CMV-seropositive group relative to the CMV-seronegative group by both raw and functional diversity, for all weighting parameters (Fig. 6a). However, combining raw with functional diversity facilitated identification of two subgroups among the subjects with known CMV status: subjects with a high raw Berger-Parker Index (^∞^*D*) were almost always CMV-seronegative (Fig. 6b), whereas subjects with low functional Berger-Parker Index (^∞^*D*_*s*_) were almost always CMV-seropositive (Fig. 6c). The reverse—low ^∞^*D* or high ^∞^*D*_*s*_—did not distinguish between the two groups. Using both measures gave a better indication of CMV status than did either one alone (Fig. 6d). The conclusion is that CMV is unlikely in the absence of large clones/expanded lineages, as has been reported, but is likely only if the large clones/expanded lineages that are present exhibit high similarity to other clones/lineages in the repertoire, or else are indeed very large (Fig. 6e). Again, the addition of functional diversity offers insight that raw diversity alone does not.

**Figure 6:**
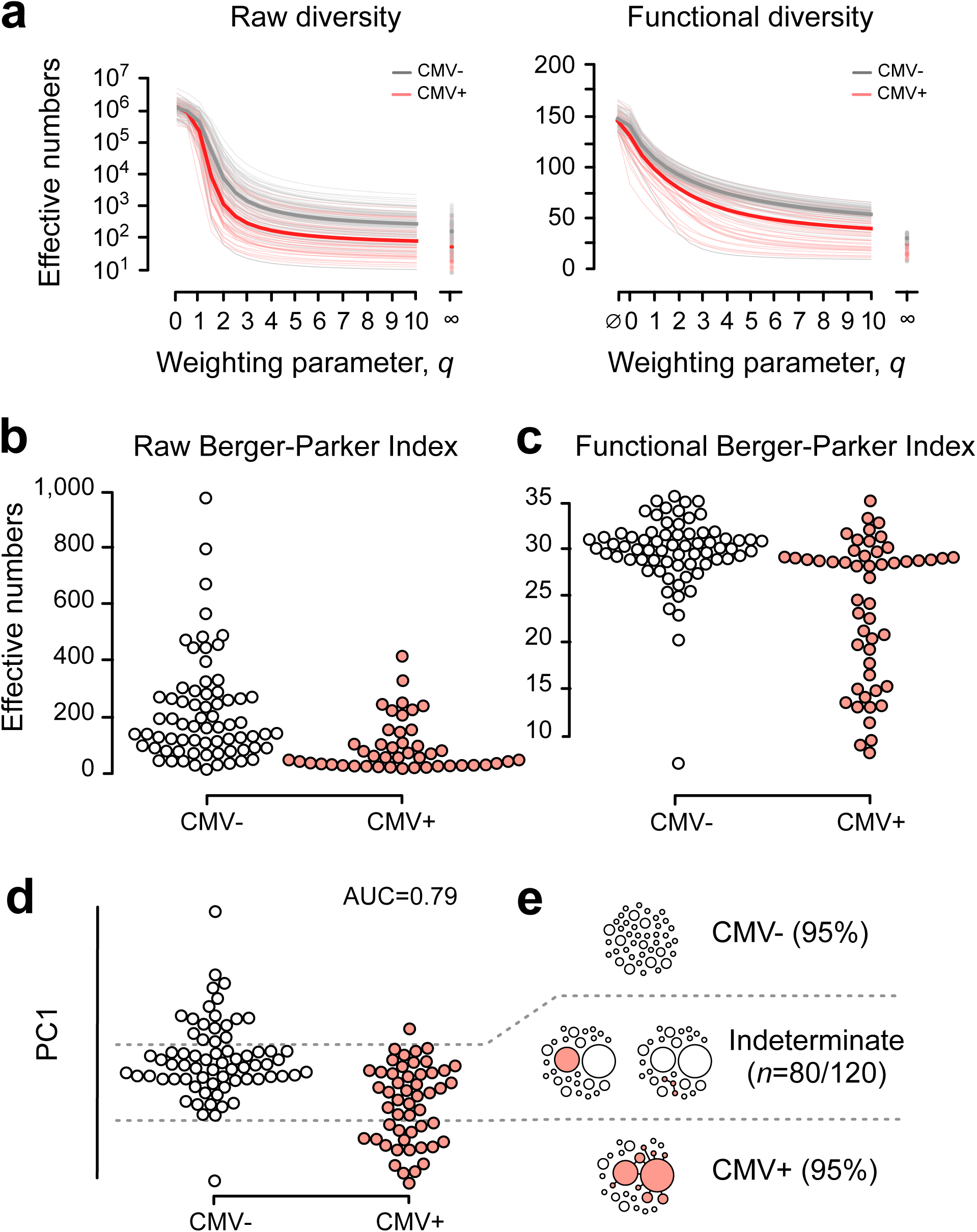
Infection. **(a)** Diversity profiles showing effective number of species as a function of weighting parameter *q* for diversity without similarity (left) and diversity with similarity (right) showing a trend toward lower diversity in CMV-seropositive individuals (red) relative to CMV-seronegative individuals (black), especially for large *q*. **(b)** Raw Berger-Parker Index (*q*=∞), which measures the largest clones, showing that high diversity—an absence of large clones—is rare in CMV-seropositive individuals. **(c)** Functional Berger-Parker Index, showing that low diversity—the presence of large clones with similarity to other clones in the repertoire—is rare in CMV-seronegative individuals. **(d)** Combining raw and functional Berger-Parker Indices (first principal component of PCA, which explains 72 percent of variance) illustrates both of the trends in (c): for the third of subjects beyond the cutoffs indicated by the horizontal dashed lines CMV serological status is assigned with an accuracy of 95 percent. **(e)** Schematic representation of the three classes revealed by combining diversity with and without similarity. Each circle is a clone; each collection of clones is a representative repertoire. Top: subjects without large clones are almost always CMV seronegative. Bottom: subjects with large clones that are similar to other clones in the sample (shown in red) are almost always CMV seropositive. Middle: repertoires with large clones that are not similar to other clones in the repertoire may be either CMV seropositive or CMV seronegative. Receiver-operator characteristic (ROC) analysis gave an area under the curve (AUC) of 0.79.

### Flu vaccination

Vaccination with a seasonal trivalent influenza vaccine (TIV) triggers clonal expansion in B cells. Previous work on five vaccinees showed likely flu-specific memory IgG lineages emerging by day 7 post-vaccination^15^. We found that combining raw and functional diversity reveals a signature of clonal expansion and selection without the need for lineage tracking (Fig. 7). We measured raw and functional diversity for IgM and IgG at day 0 (pre-administration) and day 7 from all 14 vaccinees in Vollmers’ dataset. We found that for most subjects, for IgG, raw species richness rose from day 0 to day 7 while functional species richness fell (Fig. 7a-b). This means that even as the number of sequences increased, many of the new sequences were similar to each other (or to existing sequences), and they tended to replace different-looking sequences. Meanwhile, there was no obvious pattern in IgM (Fig. 7c). Together, these results are what we would expect from clonal expansion and selection in a memory response, and thus represent a repertoire-level signature of these phenomena. Interestingly, in most cases, raw and functional entropy both fell (Fig. 7a-b, right panels). This suggests that most of the new sequences at day 7 were rare, while at the same time a subset of sequences and functional clusters grew. Thus overall, the addition of functional diversity reveals a key feature of clonal dynamics, which is not evident from raw diversity alone.

**Figure 7:**
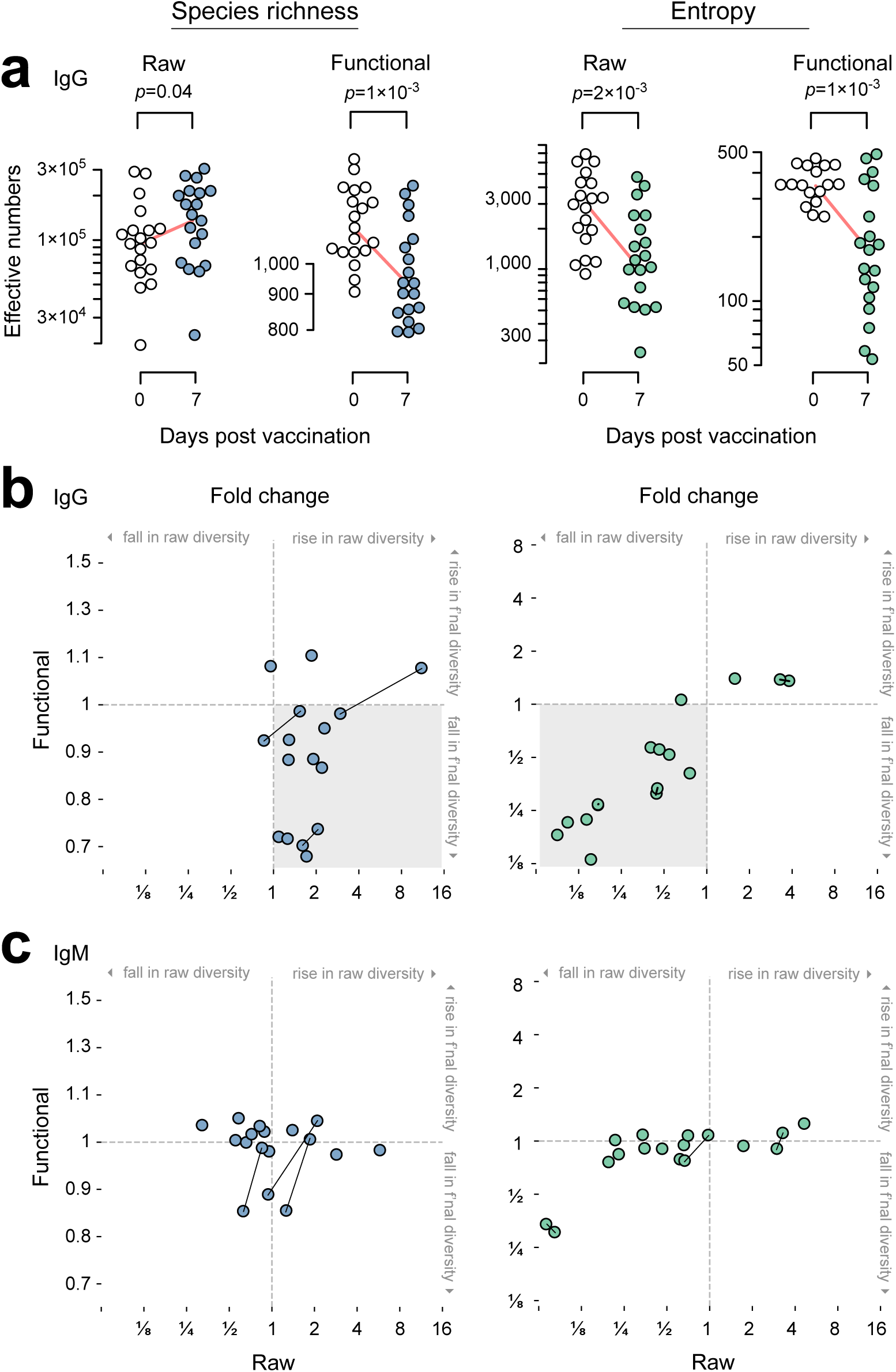
Vaccination. Raw and functional diversity together reveal clonal expansion and selection without needing lineage analysis. **(a)** In the IgG compartment, raw species richness rises while functional species richness falls in most vaccinees (left). Meanwhile raw and functional entropy both fall (right). The difference vs. species richness suggests most new sequences at day 7 are rare. (b) Meanwhile, the IgM compartment changes less by these measures.

### Aging

To explore the effect of age, we measured raw and functional diversity for TCRβ CDR3 repertoires from 41 healthy individuals aged 6-90 years old^29^ (Fig. 8). We found that raw diversity falls with age regardless of frequency parameter; a fall in only raw species richness had been reported previously^29^. Functional diversity also fell, regardless of frequency parameter. However, for species richness, four septuagenarians bucked the trend (Fig. 8, arrows): even as their raw species richness was unremarkable relative to that of other individuals of similar age, their functional species richness was similar to that of children. Only one of these four had a high likelihood of being CMV-negative; the probability that all four were CMV-negative was low. We therefore consider CMV unlikely as an explanation for their high functional species richness. Unlike their peers, these four individuals appear to have retained functional diversity among their rarest sequences. (An alternative hypothesis is that these four saw a rise in functional species richness from a lower level earlier in their adulthood, but we consider this unlikely given the overall downward trend across individuals.) We considered but excluded PCR/sequencing artifacts as the cause, as we expected such artifacts would have led to larger raw species richness, which was not observed. Thus, functional diversity can identify for further study individuals who are unremarkable by raw diversity alone.

**Figure 8:**
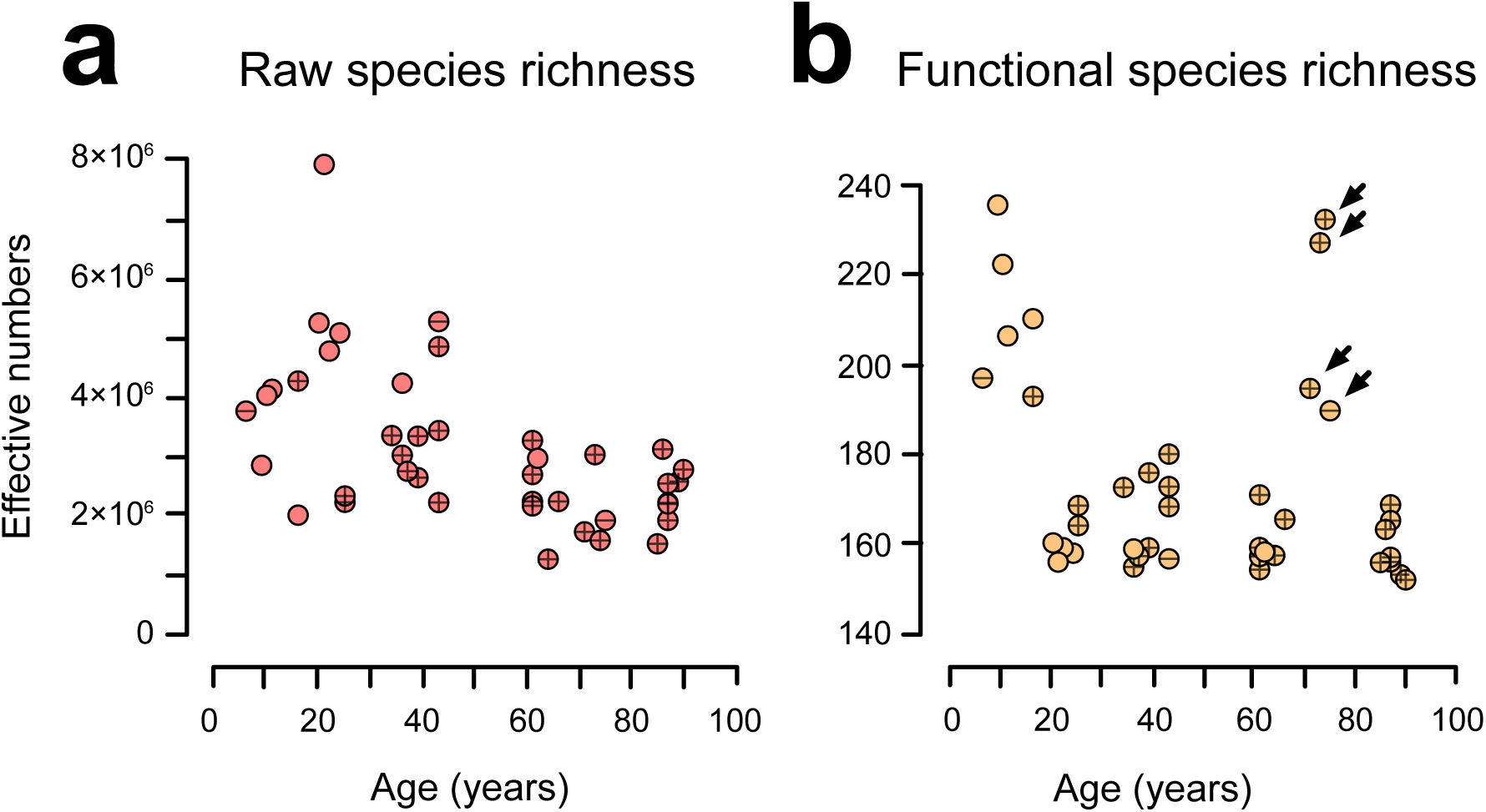
Aging. Raw and functional species richness (*q*=0) for TCRβ CDR3 repertoires from 41 healthy individuals. Arrows denote four septuagenarians who bucked the trend of lower functional species richness with age. Note that for each individual, the raw species richness is ~10-fold higher than previously reported (Britanova 2014), likely because the method we used to correct for missing species (Recon) is more sensitive than the method used in the previous report (Fisher).

## Discussion

Diversity both affects function and reflects it. In the adaptive immune system, the defining tradeoff is breadth vs. depth: a repertoire must be sufficiently diverse to contain sequences that can recognize a given target and lead to useful clones, but not so diverse that cells that express such sequences are too rare to encounter their target on biologically meaningful timescales^30,31^. To monitor immunological diversity, either diagnostically or therapeutically, it must be measured, and to measure it, must be defined. The literature increasingly recognizes that any reasonable definition of immunological diversity must account for differences in species frequency. Here we add that such a definition must also account for species’ pairwise similarity, and present a continuous framework showing that binding similarity, which leads to what we call functional diversity, provides useful insights into repertoire function.

Generically, pairwise similarity *S*_*ij*_ can be seen as governed by a tunable parameter that helps define the similarity matrix *S*, analogous to how *q* is a tunable parameter that governs the effect of differences in frequency^32^. In our study, the similarity matrix is defined by the average single amino-acid change in *K*_*d*_, an average based on over 1,300 independent measurements and multiplicative independence. Our source data is not systematic, but to our knowledge is the best available. While the present study is to our knowledge the first of its kind, it follows a long tradition of attempts to estimate the number of binding targets that can be recognized by the adaptive immune system. In these previous studies, typically a sample of B or T cells was diluted until binding to/protection from a given target was abolished, using whatever thresholds the various investigators deemed appropriate. If the limiting frequency for binding/protection was found to be, for example, 1:3,000, the conclusion was that the repertoire could recognize 3,000 different targets. This conclusion was based on the assumption that on average, all targets behave the same as the one under study. Such studies gave a wide range of functional diversities: 100 to 100,000 targets for various T-cell populations and for B cells after antigen exposure^33,34^. This range may reflect real differences in the frequencies of cells that are specific for different targets, variability in stringency or experimental setup, or some combination. Interestingly, this “how-many-can-fit” logic seems not to have been employed when testing so-called natural antibodies, which bind many targets at low affinity^35–37^. For example, one in five natural antibodies bind insulin^38^, but this is not taken to mean that the repertoire could recognize only five targets, because of presumed overlapping specificity of these antibodies for other targets. Meanwhile, theoretical studies have suggested a need for ≤10,000 binding targets, and fewer for T cells than B cells, because of major histocompatibility complex (MHC) restriction^31,39^.

In the present study, for raw diversity, we found that a typical repertoire contains on the order of 10 million unique CDR3s, well above the upper end of the range for previous estimates of the number of binding targets. The present findings are in line with other recent estimates that were likewise based on a combination of deep sequencing and statistical correction^5,11,29^. On average, the observed raw species richnesses mean that each of 100-100,000 putative binding targets can be bound by 100-100,000 unique CDR3s (10^7^/10^5^-10^7^/10^2^). From a medical perspective, such redundancy is good for treatment, because it supports the prevailing view that there are many ways to design an antibody-or TCR-based drug that will recognize a given target, but potentially a complicating factor for attempts to diagnose specific diseases based on repertoire sequence, because it suggests that signatures of exposure to a given target may often (though not necessarily always) be quite variable.

A key new finding in the present work is that functional diversity is much lower than raw diversity: repertoires contain only a few hundred functional clusters for TCRβ CDR3s and at most a few thousand for IgH. The fact that functional diversity is based on *K*_*d*_s suggests that functional diversity should correlate with the number of structurally unique, non-overlapping target clusters that CDR3s can recognize (Fig. 1). Yet the present measurements of functional species richness lie at the low end of the range of past estimates. We propose two not-mutually-exclusive explanations. First, our measurements are limited to CDR3s; variability in the rest of the antibody or TCR protein likely add to the total number of potential binding targets. This possibility is testable by extending our method to more or indeed all of the antibody or TCR sequence. Second, functional diversity may be providing a less detailed description of shape space than limiting-dilution studies: i.e., functional diversity may be a coarse graining of the target-binding landscape^40,41^ (Fig. 9). A pair of sequences may be similar enough to lie near each other in shape space, but only one may bind a given target above a threshold level of specificity in a binding study. In short, binding studies may be counting peaks while functional diversity of CDR3s counts mountains. If true, our results suggest that the landscape of TCRβ CDR3 binding is more clustered than that of IgH, such that there are on average several times as many functional IgH clusters as TCRβ. This prediction is testable through large-scale systematic binding assays to measure *K*_*d*_, or by measuring binding as a binary outcome at multiple stringency thresholds^42^. To our knowledge this is the first attempt a quantitative summary of this landscape using data from large-scale binding studies and high-throughput repertoire sequencing.

**Figure 9:**
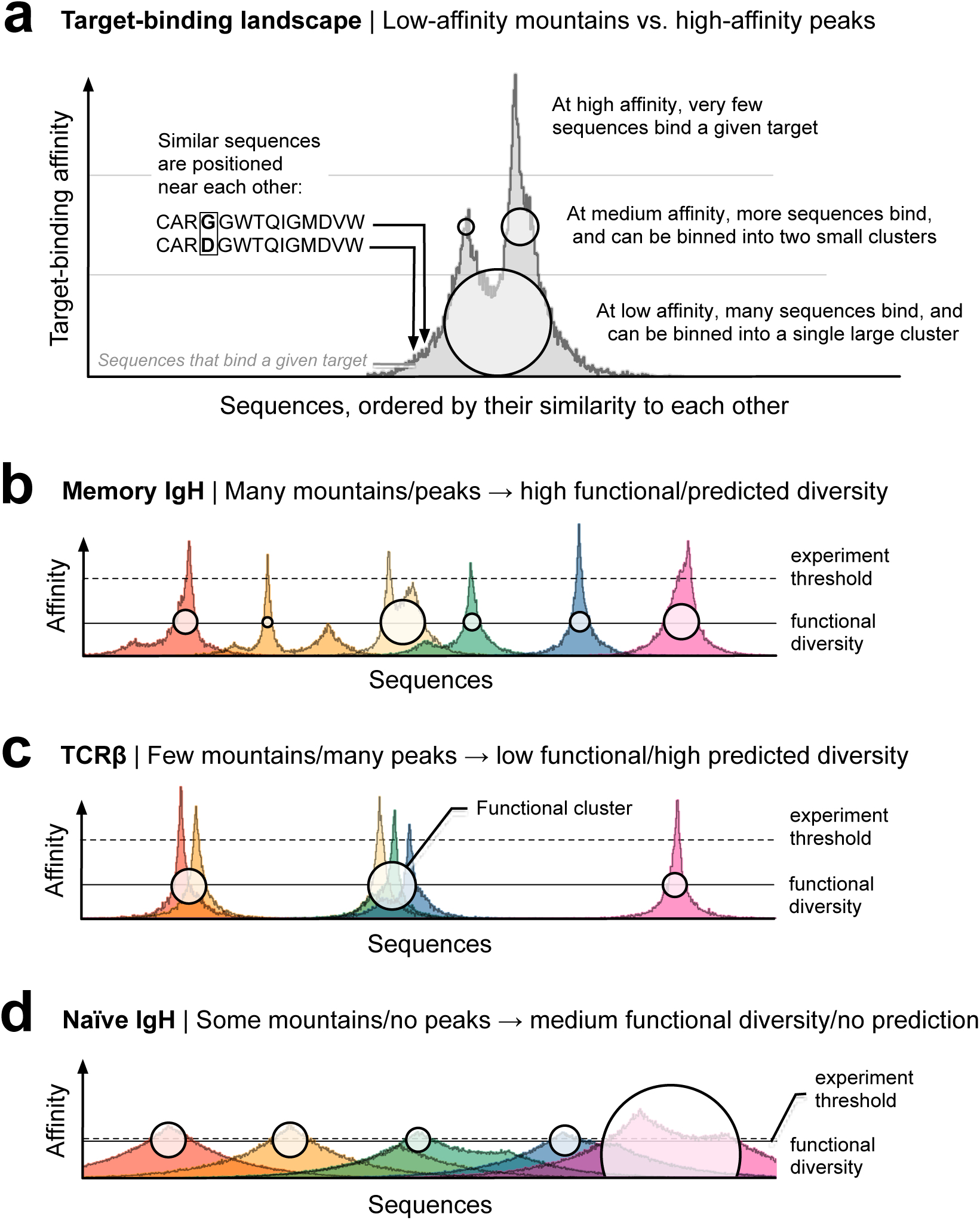
Binding landscape. **(a)** Schematic of the target-binding landscape. The gray distribution represents CDR3 sequences that bind a given target. Sequences are ordered by their similarity to each other. (In reality, similarity is a multidimensional property that makes it impossible to order sequences in a single dimension as shown here; this is done for illustrative purposes only.) The height at each sequence denotes the affinity with which it binds the target (vertical axis; measured e.g. by *K*_*d*_). Many more sequences bind the target at low affinity than at high affinity, resulting in a “mountain-and-peak” appearance. This schematic is useful for interpreting functional diversity as described in this study, and the raw diversity estimates based on previous binding studies, as described in the Discussion. (Note that in this schematic, the raw diversity as measured simply corresponds to the total number of sequences along, i.e. the width of, the horizontal axis.) At high affinity, very few sequences bind a given target. At medium affinity, more sequences bind, and can be binned into two small clusters, represented by the small circles. At low affinity, many sequences bind, and can be binned into a single large cluster, represented by the large circle. In **(b)**-**(d)**, many targets are shown. Each color corresponds to a different target; nearby targets are structurally similar. As in (a), each colored area denotes the sequences that bind a given target, as a function of binding affinity (vertical axis). Experiments usually detect the highest-affinity sequences: the peaks of the landscape (above the horizontal dotted line). The narrower the peak when it crosses the experimental threshold, the rarer specific sequences are, and the larger the number of targets that the repertoire will be estimated to bind. (For example, if 100,000 sequences are shown across the horizontal axis in each plot, and only one crosses the experimental threshold for a given target, the frequency of sequences specific for that target is 1:100,000, and the conclusion will be that there must be 100,000 such targets that the repertoire can bind. If 100 cross the experimental threshold, the conclusion will be that the repertoire can bind only 1,000 targets.) Functional diversity measures the overall contours of the landscape. Conceptually, this can be thought of as measuring the number and size of the “mountains” at a lower affinity threshold (horizontal solid lines). The differences in functional diversity between (b) memory IgH, (c) TCRβ, and (d) naïve IgH correspond to different landscapes. The raw species richness of memory IgH and TCRβ are comparable, represented here by the same width of all the plots. In addition, a similar number of sequences per target cross the experimental threshold, so estimates of the total number of targets that the repertoires can bind will also be comparable. However, less low-affinity overlap between the targets of the IgH sequences in (b) gives it higher functional species richness than the TCRβ repertoire in (c): here, six functional clusters (white circles) vs. three. (The sizes of the clusters are related to frequency-weighted functional diversity measures, i.e. larger *q*.) The sequences in the naïve IgH repertoire in (d) have only low affinity for the six colored targets, and many recognize more than one target (overlapping colored areas). Note the lower experimental threshold (horizontal dotted line), consistent with the ~10% or more of antibodies that recognize a target and the high degree of cross-reactivity in studies of natural antibodies ^39,40^. The functional diversity threshold controlled by the average effect on *K*_*d*_ of a single-amino-acid change in CDR3. If the effect were larger—or if it were amplified by e.g. raising it to a power when building the similarity matrix—the threshold would be higher, and vice versa.

Why is CDR3 functional diversity higher for IgH than for TCRβ? We hypothesize that it is for the same three reasons that there is more sequence diversity for IgH than TCRβ. First, humans have 23 D_H_ gene segments vs. only 2 D_β_ segments (a 10-fold difference), and D is the largest germline contributor to CDR3. V and J segments tend to directly contribute little more than the canonical starts and ends of CDR3s, and there are roughly similar numbers of V-J combinations in IgH as TCRβ (49×6=294 and 48×13=624, respectively; i.e. a twofold difference in the opposite direction, or an aggregate five-fold difference in favor of IgH). Second is somatic hypermutation, which diversifies IgH but not TCRβ. And third, IgH CDR3s are longer than TCRβ CDR3s, allowing for a larger number of possible sequences. Further analysis will be needed to test these possible contributions.

We have shown how functional diversity complements raw diversity to offer insight into the difference between naïve and memory repertoires, to aid in identification of disease states, and to illustrate clonal selection and other repertoire dynamics. We hope these examples will encourage others to use and/or expand our framework to investigate repertoire dynamics in other conditions, in other subsets, in the other chains (TCRα and IgL), and in other model systems such as zebrafish^23^ and mouse^3,14^. We draw attention to the fascinating difference between the number of unique sequences, which ran into the millions in most of the repertoires we investigated, and the much smaller numbers of what we call functional clusters (the effective numbers of functional diversity). The result is a “functional degeneracy” among sequences that are organized into functional clusters. Characterizing these clusters is an interesting topic for future work.

Another key finding is the perspective that functional similarity offers on diversity across individuals and on similarities and differences between people. To show that functional diversity is robust to sampling, we generated “meta-repertoires” by pooling sequences from scores and in some cases over a hundred people, including people from different ethnic backgrounds. Surprisingly, and somewhat unexpectedly given the low sequence overlap among individuals, the functional diversity of these meta-repertoires never exceeded the functional diversity of any given repertoire by more than a few fold; moreover, the functional diversity trended toward saturating in samples of just a million sequences (Fig. 3). Together, these findings indicate that any two individuals share a majority of their functional clusters, in stark contrast to the vanishingly small fraction of sequences they share. Further, these findings imply that the functional diversity of the entire population is only a few hundred clusters for TCRβ CDR3s and a few thousand for IgH, and moreover that these clusters can be sampled exhaustively by sequencing fewer than 20 individuals. It will be fascinating to test this finding by generalizing beyond the ~100 individuals studied in this experiment, including with additional ethnically and geographically diverse populations, to further examine our prediction that, contrary to conventional wisdom, the functional limits of the adaptive immune system for the entire human species are in a practical sense both finite and within reach.

Finally, the focus of this study was binding similarity, but we expect that the utility of the diversity-with-similarity framework will extend to other facets of immunology (using other similarity measures to study, e.g., somatic hypermutation) and to other fields, most readily metagenomics, sociology, oncology, and cellular cartography^43–47^. We hope this study will serve as a template for the methodology and value of incorporating similarity into the study of these and other complex systems.

## Acknowledgements

This research was supported by grants from the National Institutes of Health (NIAID K08AI11495801) and the American Heart Association (15GPSPG23830004), contracts HHSN268201500003I, N01-HC-95159, N01-HC-95160, N01-HC-95161, N01-HC-95162, N01-HC-95163, N01-HC-95164, N01-HC-95165, N01-HC-95166, N01-HC-95167, N01-HC-95168 and N01-HC-95169 from the National Heart, Lung, and Blood Institute, and by grants UL1-TR-000040, UL1-TR-001079, and UL1-TR-001420 from NCATS, as well as support from the Extreme Science and Engineering Discovery Environment (XSEDE), which is supported by National Science Foundation grant number ACI-1548562. The authors thank the investigators, staff, and participants of the MESA study for their valuable contributions (a full list of participating MESA investigators and institutions can be found at http://www.mesa-nhlbi.org), as well as the Research Computing Group at the High Performance Computing Cluster at Harvard Medical School. Finally, the authors would like to thank Drs. Mohammed Al-Quraishi and Jeremy Gunawardena for helpful conversations and comments on the manuscript.

## Methods

### High-throughput repertoires

We obtained 391 quantitative high-throughput IgH and TCRβ repertoires from 202 human subjects. These included IgH from naïve and memory B cells from DNA (*n*=3 individuals)^5^; TCRβ chains from DNA from healthy subjects known to be serologically negative for cytomegalovirus (CMV) (*n*=69 individuals)^26^ and from healthy subjects whose CMV serostatus was unknown (*n*=41 individuals)^29^; pooled barcoded IgG and IgM heavy chains from mRNA from healthy subjects before and seven days after administration of one of two influenza vaccines (*n*=28 individuals)^15^; quantitative pooled TCRβ chains from DNA for subjects who were otherwise healthy but serologically CMV positive (*n*=51 individuals)^26^; and IgH chains from DNA for subjects enrolled in the Multi-Ethnic study of Atherosclerosis (MESA; *n*=41 individuals)^25^. CDR3 annotation was performed using our in-house pipeline as previously reported^14^ and standard tools (e.g. IMGT). Details for obtaining these datasets are available from the primary publications referenced above.

### Similarity measures

A functional measure of similarity between polypeptides is how well they bind the same target. We were interested in similarity as a function of the number of amino acid substitutions (i.e., as a function of edit distance). The effect of substitutions on binding is complex and depends on the position and identity of the specific amino acids involved; many substitutions may have little or no effect, while a few may abolish binding entirely^48^. When comprehensive data are available, detailed statistical models can offer reasonable predictions of the effect of specific amino-acid substitutions^49–51^. However, this type of data does not yet exist across entire antibody and TCR repertoires, and so simpler models are required. These models are not expected to precisely predict the effects of specific substitutions, but should accurately reflect the effects of substitutions when averaged over many pairs of proteins, such as the millions of pairs in megacell-scale repertoires.

To develop a model for our similarity measure, *s*, we downloaded SKEMPI 2.0, which is to our knowledge the largest and best-curated database of experimentally measured effects of amino-acid substitution on protein-protein binding^20^. Each entry includes a Protein Data Bank (PDB) identifier^52^, the type of structural region^53^ that contains the substitution(s), one or more PDB coordinates, and (in nearly all cases) the dissociation constant (*K*_*d*_) of each member of the pair (referred to in the database and Fig. 2a as “wild type” and “mutant”). We extracted entries for all single amino-acid substitutions for which *K*_*d*_ for both wild type and mutant were recorded, and considered only entries that involved binding between antibody and antigen (*n*=797) or TCR and peptide/MHC (*n*=531; total *n*=1,328). Although amino-acid substitutions anywhere in a protein may affect binding, substitutions at the core of the binding interface are more likely to affect binding than substitutions elsewhere^53^. Therefore we split the data into core (*n*=584) and non-core (*n*=744) groups and analyzed the effect of substitution binding, measured as |log_10_(*K*_*d*__mut_/*K*_*d*__wt_)|, separately for each group.

As expected, the probability distributions for the two groups differed substantially from each other (Mann-Whitney U *p*-value 2.0×10^−33^). Substitution of a core residue had a 13-fold (geometric) mean effect on binding, consistent with prior reports^54^, while substitution of a non-core residue had a 4-fold effect. Both probability distributions were long-tailed, and were reasonably well described by exponential probability-density functions (i.e., of the form *k*e^−*kx*^). We confirmed that the distributions for antibody-antigen core residues (*n*=352) and TCR-peptide/MHC core residues (*n*=232) were similar to each other, that the distributions for antibody-antigen non-core residues (*n*=445) and TCR-peptide/MHC non-core residues (*n*=299) were also similar to each other, that within each of the antibody-antigen and TCR-peptide/MHC subgroups the distributions for core and non-core residues were different, and that these results held separately for human and non-human (nearly all of which were mouse) sequences (using the Structural Antibody Database^55^ and the Structural TCR Database^56^ to assign species), all using Mann-Whitney U and visualized as histograms. Using PyMol v2.2.0^57^, we next manually reviewed nine structures containing substitutions in human IgH or TCRβ CDR3s (1BD2, 1OGA, 3BN9, 3QDJ, 3SE8, 3SE9, 4I77, 5C6T, 5E9D) and found that to a good approximation, a constant fraction 0.15±0.05 of CDR3 amino acids consist of core residues, with no obvious difference between chain types. To estimate the effect of a single amino-acid substitution in a CDR3 in our datasets, we therefore combined core and non-core distributions with a weighting of 0.15:0.85.

The resulting distribution was again long-tailed, with most substitutions having small effects and a few having effects of many orders of magnitude (Fig. 2a). There were small spikes in the tail for substitutions with ≳60-fold effects, i.e. |log_10_(*K*_*d*__mut_/*K*_*d*__wt_)|≳1.8. A review of sources cited by SKEMPI suggested that these spikes likely reflect ascertainment bias: selective experimentation on amino acids with unusually strong effects (e.g. refs 58,59). To counteract such bias, we therefore built a high-confidence dataset using 1.8 as the cutoff. Ascertainment bias in two- and three-amino-acid substitutions is expected to follow the square and cube of the bias in single-amino-acid substitutions, respectively, precluding rigorous conclusions from being drawn from independence testing. However, comparison with those groups was broadly consistent with either multiplicative (*s*=*c^m^*, where *s*=similarity; *c*=the cost of binding, i.e., 1/(fold effect); *m*=edit distance) or additive (*s*=*c/m* for *m*≥1) independence. Because additive effects result in higher pairwise similarities and therefore smaller repertoire diversities than multiplicative effects, the multiplicative-independence model is more conservative for studying the effects of similarity on diversity. We therefore chose the multiplicative model for further analysis.

To determine the similarity between two CDR3s with edit distance *m*, we sampled independently from the high-confidence dataset *m* times, and multiplied the costs together. We confirmed that on average, the results of this stochastic sampling were the same as deterministic calculation of *s*=*c*^*m*^ with *c*≈0.55. We performed sensitivity analysis based on lower-confidence cutoffs (down to *c*=0.48) and alternative assumptions (up to *c*=0.60). This resulted in somewhat higher or lower diversity values, but qualitative patterns were robust to these perturbations.

### Diversity measures

We calculated ^*q*^*D* according to Eq. (1) as previously described^9,11^ and ^*q*^*D*_*s*_ according to Leinster and Cobbold^16^ following Eqs. (2)-(3). We corrected ^*q*^D for sampling error using Recon (github.com/ArnaoutLab/Recon; default settings) as previously described^11^. We note Hill’s framework^9^ (Eq. (1)) has inspired several methods for incorporating similarity into diversity measurements, each of which retains useful features of Hill’s framework^16,32,60,61^. Two of the new frameworks were introduced with explicit discussion of how to decompose population-level diversity into within- and between-group components^16,60^. Each of these has advantages and disadvantages over the other ^62^. We chose Leinster and Cobbold’s framework here because we found it easier to apply and interpret. For readability, we made two minor changes to the notation, from ^*q*^*D*^***Z***^ to ^*q*^*D*_*s*_ and from **Zp**_*i*_ to ***S**_i_*.

Use of this framework raised two issues that we addressed. First, its *q*=0 measure, ^0^*D*_*s*_, depends on frequency albeit to a very small extent, unlike the Hill framework’s *q*=0 measure, ^0^*D*, which is species richness (and is independent of frequency). Therefore, as a more direct comparison to species richness, we calculated ^0^*D*_*s*_ both with frequency information and without it (i.e., setting the frequencies of each of the *n* species to 1/*n*). We refer to the latter as ^0^*D*_*s*_ (“*D*-null”). Second, it has been shown that this framework can result in unreasonably low diversity values when most of the off-diagonal entries of the similarity matrix are far from zero, resulting in an insensitivity to *q* ^67,69^. We expected most of our off diagonals to be close to zero, since our similarity measures directly or indirectly involve exponential decay, which generates small values, but confirmed that most of the off diagonals in our similarity matrices were indeed close to zero by plotting histograms. Consequently, our measures were sensitive to *q*, as desired and expected (Figs. 5-8).

### Robustness analyses

For robustness analyses, IgH and TCRβ were analyzed separately. The upper-bound/worst-case scenario for IgH was evaluated by constructing a “meta-repertoire” by combining IgG sequences of subjects before vaccination (n=28 individuals)^15^, sequences from memory cells from healthy subjects from public database (*n*=3 individuals)^5^, and sequences from subjects enrolled in MESA study (*n*=41 individuals)^25^, and sampling from this meta-repertoire without regard to the frequency of sequences. We chose IgG/memory sequences where possible because those sets exhibited higher functional diversity than naïve sets, and we were interested in maximizing diversity. We ignored the frequency of sequences for the same reason: uniform frequency maximizes diversity, other things equal. For TCRβ, we constructed a meta-repertoire by combining sequences from CMV seronegative individuals (*n*=69 individuals)^26^ and again sampling at uniform frequency. We chose CMV seronegative individuals for the same reason as we chose memory/IgG sequences above: seronegative individuals exhibited higher diversity. For both IgH and TCRβ, including all sequences lowered diversities slightly. The representative samples were from subject D3 for IgH (from DNA), subject SRR960344 for IgH (from mRNA), and subject Keck0070 for TCRβ (CMV seronegative). CDR3 sequences were sampled proportional to their frequency in the repertoire.

